# Terminal neuron localization to the upper cortical plate is controlled by the transcription factor NEUROD2

**DOI:** 10.1101/651273

**Authors:** Gizem Guzelsoy, Cansu Akkaya, Dila Atak, Cory D. Dunn, Alkan Kabakcioglu, Nurhan Ozlu, Gulayse Ince-Dunn

**Affiliations:** Department of Molecular Biology and Genetics, Koç University, Istanbul, Turkey; Stem Cells and Metabolism Research Program, Faculty of Medicine, University of Helsinki, Helsinki, Finland; Institute of Biotechnology, Helsinki Institute of Life Science, University of Helsinki, Helsinki, Finland; Physics Department, Koç University, Istanbul, Turkey; Center for Genomics and Systems Biology, Department of Biology, New York University, New York City, NY, USA; Translational Medicine Research Center, Koç University, Istanbul, Turkey

**Author notes:** These authors contributed equally to this work (Gizem Guzelsoy, Cansu Akkaya).

## Abstract

Excitatory neurons of the mammalian cerebral cortex are organized into six functional layers characterized by unique patterns of connectivity, as well as distinctive physiological and morphological properties. Cortical layers appear after a highly regulated migration process in which cells move from the deeper, proliferative zone toward the superficial layers. Importantly, defects in this radial migration process have been implicated in neurodevelopmental and psychiatric diseases. Here we report that during the final stages of migration, transcription factor Neurogenic Differentiation 2 (*Neurod2*) contributes to terminal cellular localization within the cortical plate. In mice, *in utero* knockdown of *Neurod2* results in reduced numbers of neurons localized to the uppermost region of the developing cortex, also termed the primitive cortical zone. Our ChIP-Seq and RNA-Seq analyses of genes regulated by NEUROD2 in the developing cortex identify a number of key target genes with known roles in Reelin signaling, a critical regulator of neuronal migration. Our focused analysis of regulation of the *Reln* gene, encoding the extracellular ligand REELIN, uncovers NEUROD2 binding to conserved E-box elements in multiple introns. Furthermore, we demonstrate that knockdown of NEUROD2 in primary cortical neurons results in a strong increase in *Reln* gene expression at the mRNA level, as well as a slight upregulation at the protein level. These data reveal a new role for NEUROD2 during the late stages of neuronal migration, and our analysis of its genomic targets offer new genes with potential roles in cortical lamination.

## Introduction

The cerebral cortex of higher animals is composed of numerous excitatory and inhibitory neuron types organized into six anatomical and functional layers. Developmentally, excitatory neurons originate from neural stem cells that divide in the ventricular zone of the anterior neural tube, then migrate radially to the overlying cortical plate. While neural stem cells initially divide symmetrically during an increase of the proliferative population, the division mode transitions to asymmetric division as embryogenesis progresses. One daughter cell from each cell division remains at the ventricular zone, while the other daughter exits cell cycle and initiates a complex mode of radial migration to the border between the cortical plate and the marginal zone. As development progresses, each wave of migrating neurons reaches the surface by passing the nascent cortex in an inside-out manner, eventually giving rise to the layered structure of the cerebral cortex ^1,2^. Consequently, early-born neurons populate the deep layers, and late-born neurons settle in the superficial layers. The precise control of the radial migration process is essential for the proper formation of the six cortical layers. Importantly, in mice and humans, mutations, infections or environmental agents that perturb cortical migration cause cortical structural malformations, intellectual disability, neuropsychiatric diseases, and seizures ^3-5^.

A closer examination of the radial migration process reveals a series of different modes of cellular motility, as opposed to a uniform directed movement from the ventricular zone toward the superficial marginal zone. Initially neurons are multipolar in morphology as they emerge from the ventricular zone, which is followed by a transition to a bipolar morphology and locomotion on radial glia fibers through the intermediate zone and much of the cortical plate ^6,7^. Upon approaching the border between the cortical plate and the marginal zone (MZ), they terminally translocate from the radial glia fibers. Terminal translocation is accompanied by positioning of neuronal cell bodies in the uppermost layer of the cortex, also termed as the primitive cortical zone ^8^. During this developmental period, the marginal zone is populated by Cajal-Retzius cells which secrete the extracellular glycoprotein REELIN, considered a stop signal for migrating neurons ^9^. Migrating neurons express the REELIN receptors VLDLR and LRP8 (APOER2), as well as the intracellular adaptor molecule DAB1, which conveys receptor activation to downstream signaling pathways. Ligand binding induces the endocytic internalization of REELIN and phosphorylation of DAB1 ^9^. Activation of REELIN signaling in neurons induces anchorage of nascent primary dendrites to the extracellular matrix of the marginal zone, stabilization of the neuronal skeleton, and termination of migration ^10-15^. Loss of REELIN, VLDLR, APOER2 or DAB1 function causes profound defects in cortical lamination; the cortical layers develop in an inverted pattern, with early- and late-born neurons being located in superficial and deep layers, respectively ^13,16-21^.

Individual steps of radial migration are controlled by the combinatorial expression of specific transcription factors (TFs) ^2^. Prominent among these transcription factors are the bHLH family members, including the Neurogenins and the NeuroDs. In addition to their roles in neuronal cell fate specification and neuronal differentiation, these factors have also been shown to control specific steps of cortical radial migration described above ^22-24^. A key member of the NeuroD family of neurogenic bHLH TFs is NEUROD2. The *Neurod2* gene is highly expressed in the developing cortex and its expression persists, albeit at low levels, into adulthood in cortical excitatory neurons ^25,26^. Interestingly, several recent studies by our group and others have implicated NEUROD2 in the radial migration process of cortical neurons and have provided a general overview of its downstream genetic targets ^27,28^. However, how NEUROD2 regulates the expression of its downstream targets, and how this regulation impacts cortical migration, remain largely unknown.

We previously characterized the genetic targets of NEUROD2 in the cerebral cortex at two developmental timepoints: embryonic day 14.5 (E14.5), representing the peak of neurogenesis and migration in the mouse cortex; and postnatal day 0 (P0), representing the onset of neuronal differentiation, dendritic growth and synaptogenesis ^26,27^. Here, we carry out a comparative analysis of these two datasets and overlay it with transcriptomics analysis of primary neurons where *Neurod2* expression is knocked down. We find that NEUROD2 exhibits quantitative and qualitative differences in target-selectivity at these two developmental timepoints. From our postnatal dataset, we identify several gene targets with known roles in neuronal projection development, such as the Ca^2+^/Calmodulin-dependent Kinase IV (*Camk4*) gene, and we demonstrate that suppressing *Neurod2* expression in primary cortical neurons causes dendritic differentiation defects. Using our embryonic dataset we identify numerous NEUROD2 targets with known functions in the Reelin signaling pathway, such as *Reln, Dab1* and *Lrp8*. Furthermore, suppressing *Neurod2* expression by *in utero* shRNA electroporation caused a defect in cellular positioning of neurons to the primitive cortical zone. Our results point to NEUROD2 as a regulator of the terminal stage of radial migration that acts, at least in part, by regulating genes functioning in the Reelin pathway. Our dataset also provides novel candidate genes that might have functions in different aspects of cortical radial migration. Future experiments rescuing individual target genes with roles in neuronal migration in a *Neurod2*-deficient background will uncover the exact role of how this TF orchestrates critical pathways for proper cortical lamination.

## Results

### NEUROD2 binds to overlapping and unique gene-sets in embryonic and postnatal cerebral cortex tissue

To evaluate whether NEUROD2 interactions with specific genomic sites are temporally regulated, we compared NEUROD2 binding profiles between embryonic day 14.5 (E14.5) and postnatal day 0 (P0). We retrieved binding scores from our previously published NEUROD2 ChIP-Seq datasets that represent the amount of relative TF binding to target sequences and carried out a comparative analysis using the E14.5 and P0 results ^26,27^. Based upon this comparison, we identified a large number of binding sites that were exclusively targeted at E14.5, and fewer sites that were targeted at both developmental stages or were specific to P0 (Supplemental Material 1) (Fig. 1A). For both datasets, we had previously reported that a majority of NEUROD2 binding sites were not located at promoters, but rather at distant regulatory elements. Therefore, we decided to employ a target gene prediction approach, called the Closest Gene Method, which generates a score based on the number of TF binding sites and their proximity to the transcription start site of each gene ^29^. We calculated Closest Gene scores for the P0 dataset and compared it to the E14.5 dataset which we had previously published (Supplemental Material 2) ^27^. Genes with a Closest Gene score above a threshold value were predicted as targets. Comparison of these predicted targets between two developmental stages revealed a large number of E14.5-specific genes, as well as a smaller number of target genes either only in the P0 developmental stage or at both stages (Fig 1A).

**Figure 1.**
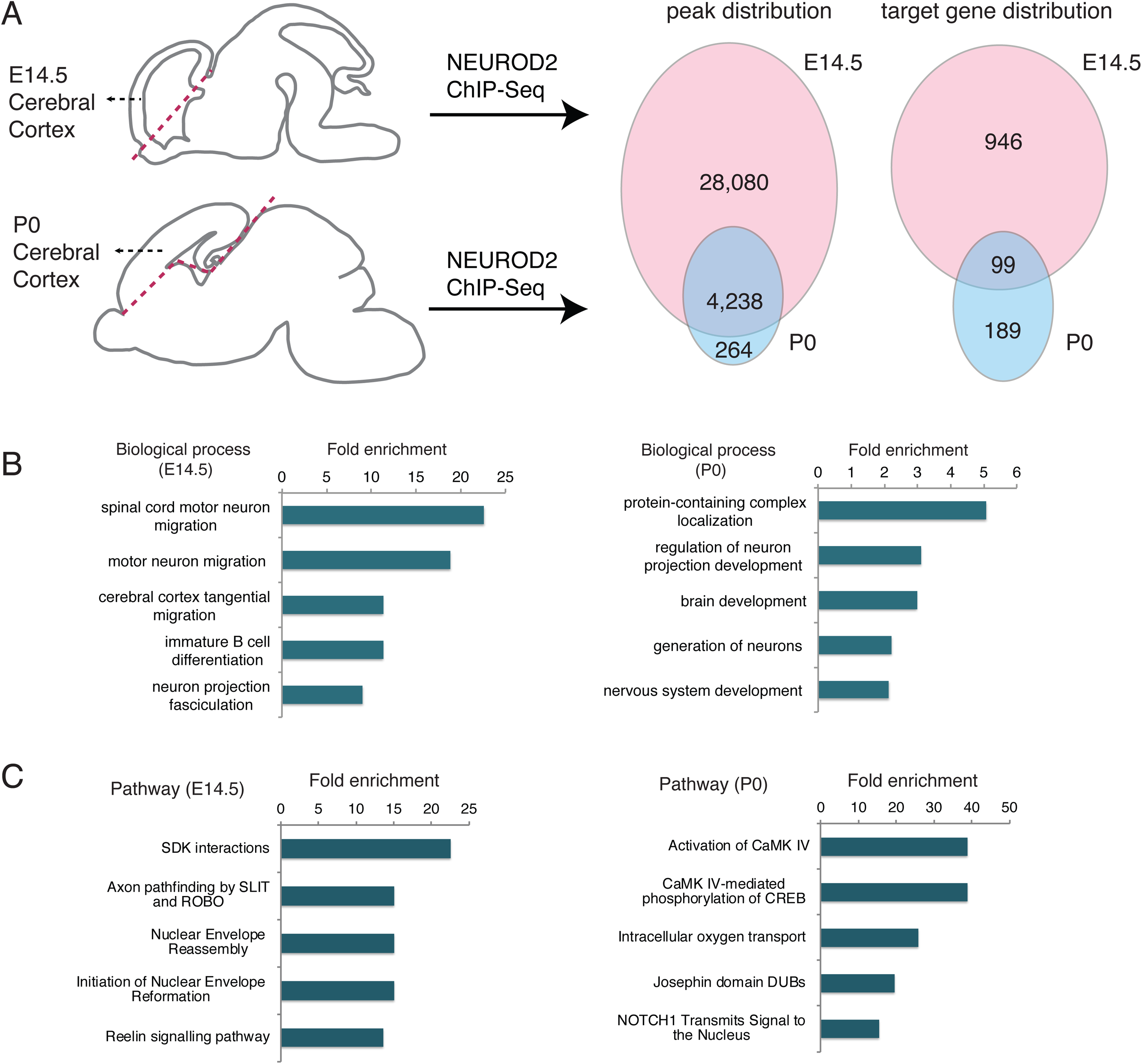
NEUROD2 binds to overlapping and unique target sites in embryonic and postnatal cerebral cortex. (A) NEUROD2 binding sites and target genes were compared between embryonic day 14.5 (E14.5) and postnatal day 0 (P0) cerebral cortical tissue. Numbers of genome-wide binding sites and target genes are based upon previously acquired ChIP-Seq data ^26,27^. Target genes were identified based on the total number of binding sites proximal to the transcription start sites of individual genes ^29^. (B) Gene ontology analysis of NEUROD2 target genes from P0 and E14.5 datasets (geneontology.org). (C) Reactome pathway analysis of NEUROD2 targets from P0 and E14.5 datasets (reactome.org).

In order to determine whether NEUROD2 gene targets at embryonic and postnatal timepoints represented functionally distinct gene groups, we carried out gene ontology analysis. Not surprisingly, at both tested developmental stages, genes related to nervous system development were significantly enriched (Fig. 1B). However, genes associated with migration of different types of neurons were specifically enriched at E14.5. To gain better insight into the signaling pathways regulated by NEUROD2, we performed pathway analysis at the two stages using the Reactome database ^30,31^. In addition to pathways whose significance is not immediately clear to us, our results also demonstrated that embryonically expressed NEUROD2 might control pathways related to cellular adhesion, synaptic connectivity, migration, and postnatal neuron projection development. Furthermore, NEUROD2 appeared to impinge upon calcium-mediated transcription, including CaMKIV transactivation of CREB (cAMP-response element binding protein) (Fig. 1C, Supplementary Material 3). Collectively these data suggested that NEUROD2, a TF that is expressed throughout the lifetime of cortical excitatory neurons, may regulate distinct functions during different developmental events.

### NEUROD2 is required for dendritic differentiation of cortical neurons

Since our analysis of our postnatal dataset suggested a role for NEUROD2 in neuronal projection development, as well as of CaMKIV-mediated transcription, a known regulator of dendritic differentiation ^32^, we asked whether neurons silenced for *Neurod2* also exhibited generalized dendritic arborization defects. Toward this aim, we cultured primary cortical neurons and transfected them with a short hairpin RNA targeting *NeuroD2* transcripts (shNeurod2-1) at low efficiency to achieve knockdown in isolated neurons (Fig. 2A). We then quantified dendritic arborization of transfected cells, also marked by EGFP expression, by Sholl analysis, an approach that reports upon the number of dendrite branches as a function of distance from the neuronal soma ^33^. Our results demonstrated that while the number of primary dendrites protruding from the soma were comparable between the two conditions, a significant reduction in higher order dendritic branches was observed in neurons where *Neurod2* expression was silenced (Fig. 2A and B). While this experiment pointed to a requirement for NEUROD2 in dendritic growth in cortical neurons, future experiments will uncover which specific target genes are functioning in dendrite arbor development.

**Figure 2.**
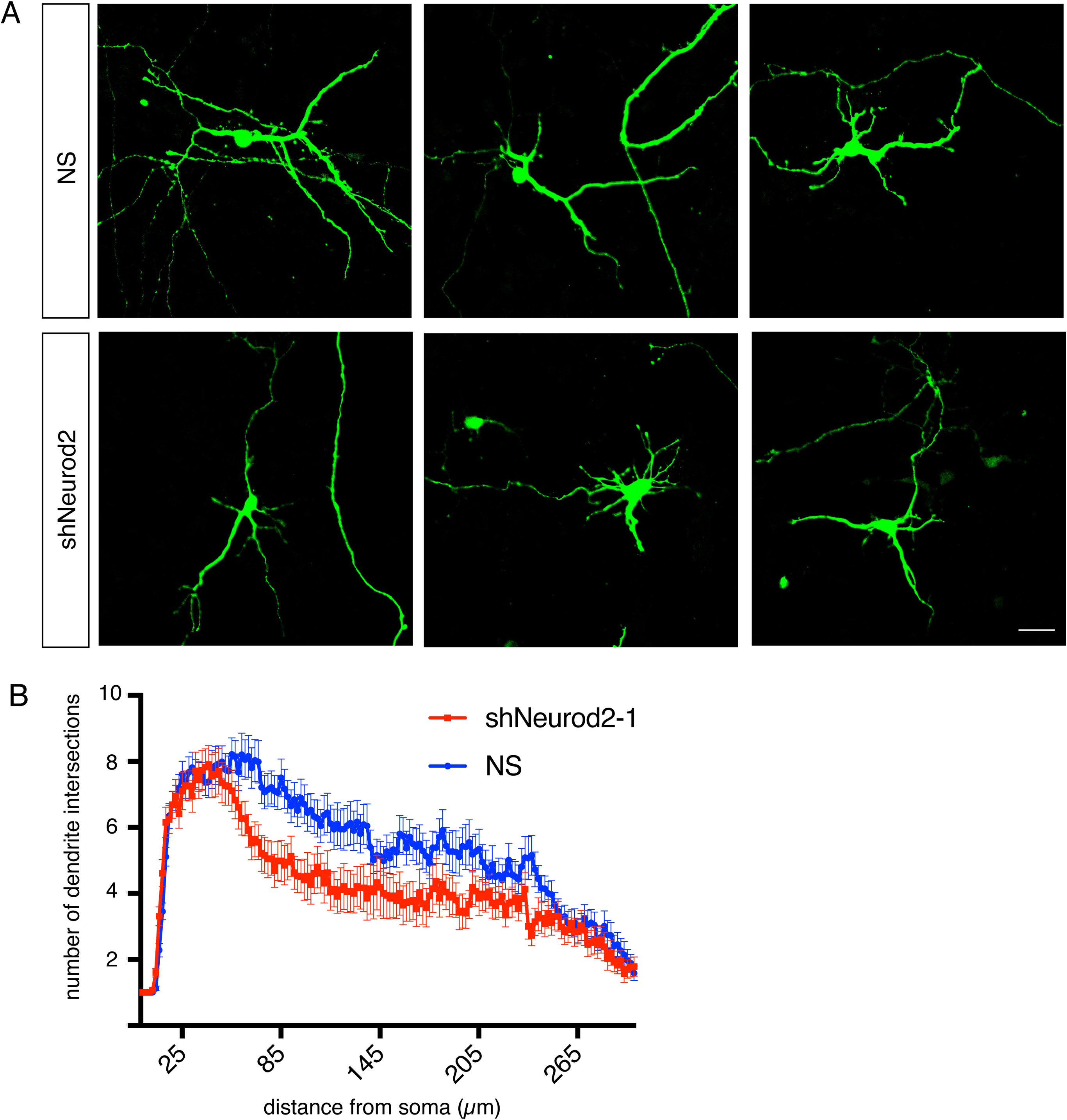
NEUROD2 is required for normal dendrite development in primary cortical neurons. (A) Primary cortical neurons from E14.5 embryos were transfected with NS (non-silencing) shRNA or shNeurod2-1 at 2 days DIV and fixed at 5 DIV. Images were captured by confocal microscopy, scale bar: 20 µm. (B) Dendrite development was quantified by Sholl analysis. n=35 neurons derived from two separate cultures. Bars indicate S.E.M. p=0.0026 by two-way ANOVA.

### RNA-Seq analysis reveals NEUROD2-regulated target genes in primary cortical neurons

To further focus our attention upon genes for which expression is regulated by NEUROD2, we silenced *Neurod2* expression in primary cortical cultures using two validated shRNAs ^26^ and analyzed gene expression changes. When gene expression was compared to cells treated with a control shRNA, we found that 25 genes were down-regulated and 9 genes were up-regulated upon silencing of *Neurod2* (Supplemental Material 4) (Fig. 3A). As expected, we detected *Neurod2* as one of the significantly downregulated genes. Genes perturbed by *Neurod2* depletion included those expressing DNA binding proteins (*Arid5a, Ddit4, Ubn2, Trim66, Zglp1*, and *Sp8*); regulators of extracellular matrix, adhesion and synaptogenesis (*Thbs1, Reln, Col3a1, Col1a1*, and *Ank1*), metabolic genes (*Shmet2, Tkt, Pck2*, and *Pde12*); signaling molecules (*Chac1* and *Sesn2*) and a translational regulator (*Rps6ka4*). Next, we asked whether any of the regulated targets were also directly bound by NEUROD2 as revealed by our ChIP-Seq data. We found that 14/34 genes harbored binding sites for NEUROD2, with the remaining (20/34) potentially represented secondary or off-target effects (Supplemental Material 4). Among the direct and regulated targets of NEUROD2, we were intrigued to identify *Reln*, a gene with essential roles in neuronal migration, dendrite development and synaptogenesis. Our previous findings had also implicated NEUROD2 in regulating the signaling pathway activated by this extracellular matrix protein ^27^.

**Figure 3.**
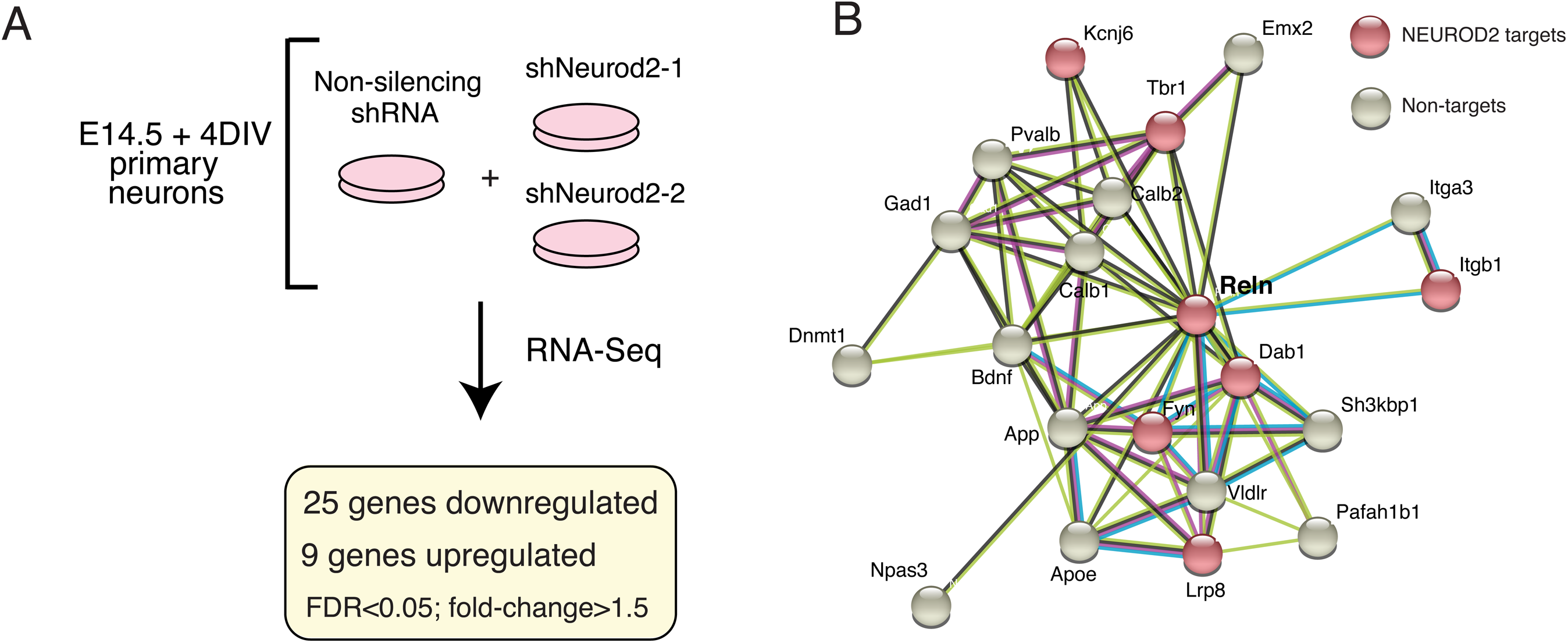
RNA-seq analysis of primary cortical neurons silenced for *Neurod2* expression. (A) RNA-Seq analysis was carried out on primary cortical neurons electroporated with a NS (non-silencing) shRNA or one of the two independent shRNAs against *Neurod2*. 25 genes were down-regulated and 9 were up-regulated genes upon knockdown of *Neurod2*. fold-change>1.5, FDR<0.05. (B) A protein interactome network of REELIN revealed additional genetic targets of NEUROD2 that also interact with REELIN. Interactome analysis was based on the String Database ^34,35^.

Next we predicted the protein interactome of REELIN using the String database ^34,35^ and asked whether additional NEUROD2 targets were found in this network. As well as the previously reported targets, REELIN receptor *Lrp8*, intracellular adaptor molecule *Dab1*, and membrane-associated tyrosine kinase *Fyn* ^27^, we found that we had also identified additional REELIN-related targets, such as *Itgb1, Tbr1* and *Kcnj6* as NEUROD2 targets that interact with REELIN (Fig. 3B). Furthermore, in order to achieve a comprehensive view of NEUROD2 targets with roles in cortical lamination, we manually searched the literature with our list of target genes and compiled a list of 116 genes with previously demonstrated roles in neuronal migration (Supplemental Material 5). These results raise the possibility that NEUROD2 might regulate cortical neuron migration via fine-tuning the expression of several downstream targets. However, given that *Reln* was the only target that displayed gene expression changes in our RNA-Seq analysis following *Neurod2* depletion, we decided to further investigate the relationship between NEUROD2 and REELIN expression.

### NEUROD2 regulates the expression of Reln gene in cortical neurons

In mice, *Reln* gene consists of 65 exons, spanning approximately 460 kb of genomic sequence ^36^. Upon inspection of NEUROD2 binding sites along this gene we identified four prominent peaks at introns 21, 22, 55 and 63 (Fig. 4A and B). Initially, we confirmed binding to all four peak sequences by ChIP followed by quantitative PCR (ChIP-qPCR) using cortical tissue from E14.5 and P0 pups (Fig. 4C). We normalized our qPCR signal individually from all four peaks to the amount of input DNA and compared to negative control samples where immunoprecipitation was carried out with an antibody against GFP. Our results suggested quantitatively more binding to all four intronic sequences from E14.5 cortex when compared to samples from P0 cortex (Fig. 4C).

**Figure 4.**
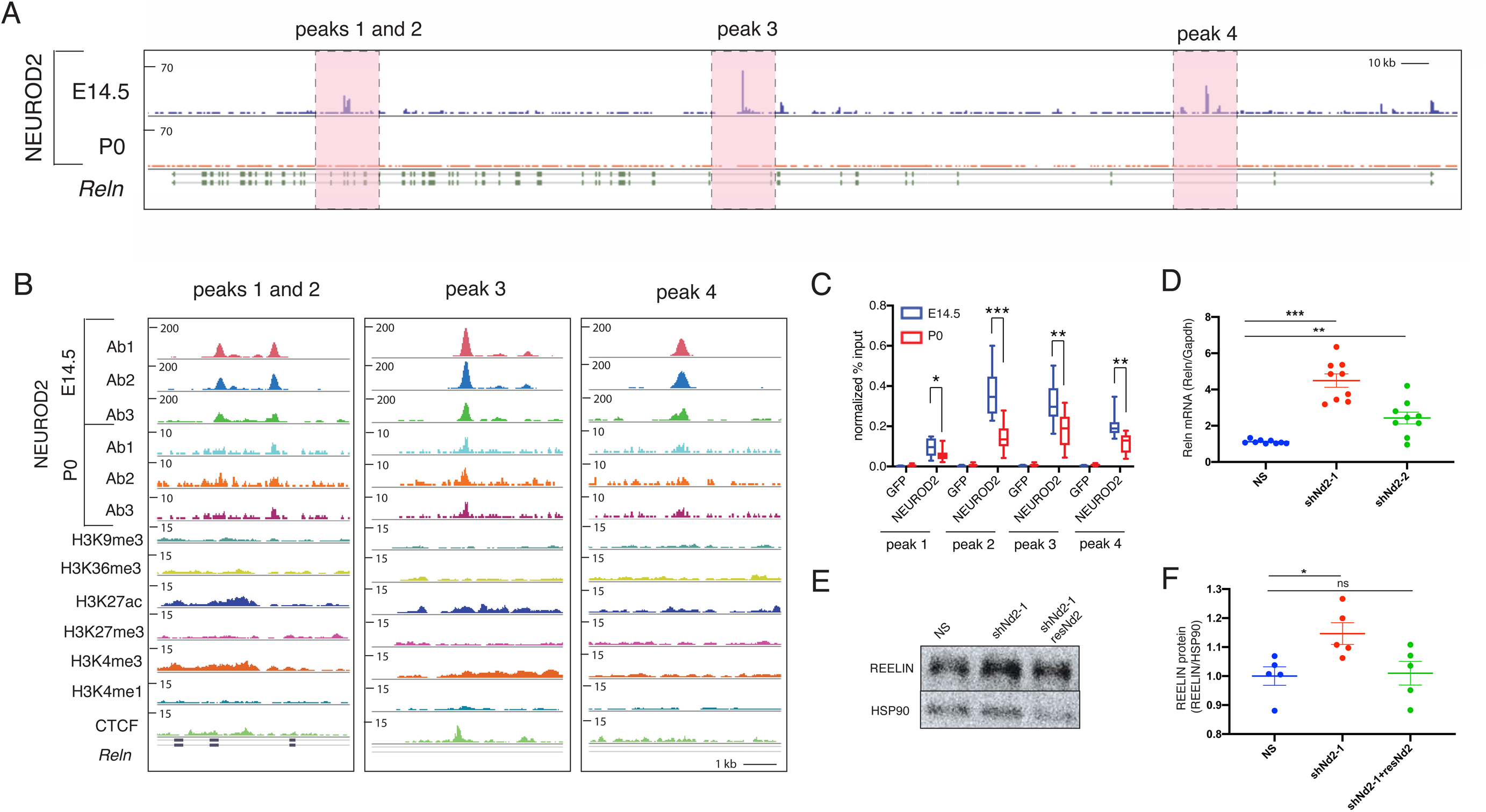
NEUROD2 binding to introns along the *Reln* gene is associated with suppression of *Reln* gene expression. (A, B) Four prominent intronic NEUROD2 binding sites are plotted on the *Reln* gene. Ab1, Ab2 and Ab3 represent ChIP experiment carried out with three separate NEUROD2 antibodies. Several different histone modifications corresponding to promoters (H3K4me3), enhancers (H3K4me1), actively transcribed (H3K27ac and H3K36me3) or repressed (H3K9me3 and H3K27me3) chromatin are not enriched in NEUROD2 binding sites. A slight enrichment of CTCF binding within intron 3 of *Reln* is detected. (C) NEUROD2 binding to all four intronic regions is confirmed by ChIP-qPCR. Immunoprecipitation with an unrelated GFP antibody is used as a negative control. n= 12 (three biological x four technical replicates). Bars represent S.E.M. *p<0.05, **p<0.005, ***p<0.0001 by two-tailed unpaired t-test. (D) RT-qPCR analysis of *Reln* gene expression reveals significant upregulation upon knockdown of *Neurod2* with two separate shRNAs. n= 9 (three biological x three technical replicates). p<0.0001 by one-way ANOVA. ***p<0.0001, **p<0.01 by post hoc Sidak multiple comparison tests. Bars represent S.E.M. (E) Immunoblotting of REELIN from 5 DIV primary cortical cultures electroporated with NS, shNeurod2-1 or with shNeurod2-1 together with a plasmid expressing an shRNA resistant Neurod2 cDNA (resNd2). The protein blot has been probed with REELIN antibody and HSP90 antibody as a loading control. Cropped parts of the gel is indicated with a line. (F) Quantification of REELIN protein amounts from five independent experiments. Data is presented after normalizing to HSP90 loading control. *p=0.02 by two-tailed unpaired t-test. Abbreviations: NS (non-silencing), shNd2-1 (shNeurod2-1), shNd2-2 (shNeurod2-2), resNd2 (rescue Neurod2), ns (non-significant).

In order to gain insight into the transcriptional activity of these sites in the embryonic cortex, we overlaid our NEUROD2 binding profiles with those representing different histone marks associated with promoters (H3K4me3), enhancers (H3K4me1), actively transcribed (H3K27ac and H3K36me3) or silenced (H3K9me3 and H3K27me3) chromatin (Fig. 4B). We did not observe a significant enrichment of either of these histone marks at NEUROD2 binding sites. Our observation might be indicative of a true lack of enrichment of specific histone marks at these loci, or alternatively, might be due to the dilution of ChIP-Seq signals of histone marks acquired from bulk cortical tissue containing many cells that do not express *Neurod2*. Interestingly, we did observe a modest enrichment of CTCF (CCCTC-binding factor) at the third peak we analyzed (Fig. 4B). This TF has previously been shown to act as a chromatin insulator and as either a transcriptional repressor or activator in a context-dependent manner ^37-39^. Hence, beyond functioning as a typical promoter-associated TF, NEUROD2 might have additional roles in formation of higher-order chromatin structures. Future experiments conducted with purified, *Neurod2*-expressing neurons will resolve how the binding of post-translationally modified histones and other factors regulating chromatin structure are localized to and regulated at NEUROD2 binding sites.

During radial migration, *Reln* is expressed and secreted by a transient group of neurons located in the marginal zone called Cajal-Retzius (CR) cells ^16,21^, which do not express *Neurod2* ^40^. On the other hand, migrating neurons express *Neurod2*, genes encoding REELIN receptors *Lrp8* (*Apoer2*) and *Vldlr*, as well as components of their downstream signaling pathway, such as the adaptor protein *Dab1*. Recent single cell RNA-Seq analysis also point to a mutually exclusive pattern of expression of *Reln* and *Neurod2* within the CR cells and within excitatory neurons of the developing cortex ^41^. Interestingly, our RNA-Seq results also suggested NEUROD2 as a negative regulator of *Reln* expression (Supplemental Material 4). To further explore the possibility that NEUROD2 represses *Reln*, we knocked down *Neurod2* by two different shRNAs in primary cortical neurons. Indeed, we observed a significant upregulation in *Reln* mRNA expression (Fig. 4D) with the more efficient shNeurod2-1 RNA ^26^ leading to a more robust upregulation in *Reln* transcript levels compared to those neurons transfected with shNeurod2-2 (Fig. 4D). Next, by immunoblotting we determined how REELIN protein levels were affected upon knockdown of *Neurod2* in our primary cortical cultures. To our surprise, we observed a much weaker (approximately 15% of the control) but significant (p=0.02) increase in REELIN at the protein level. To demonstrate the specificity of this effect, we rescued the increase in REELIN levels by co-expressing a previously characterized ^26^ shRNA-resistant *Neurod2* cDNA (Fig. 4E and F). Taken together, our results suggest that NEUROD2 strongly suppresses expression of *Reln* mRNA in cortical neurons. However, this regulation of gene expression at the mRNA level only modestly affects levels of REELIN protein.

Next we determined how REELIN protein levels were affected in the embryonic cortex upon silencing *Neurod2* expression via *in utero* electroporation of shNeurod2-1. We electroporated E14.5 embryos, fixed the cortices at E17.5 using paraformaldehyde, and carried out immunofluorescence staining against REELIN in cortical sections. In agreement with previous findings, we observed high levels of REELIN protein in the MZ, but not the CP or the SVZ-IZ of control samples (Fig. 5A and B). Our quantification of immunofluorescence signal intensity revealed no difference within the CP and SVZ-IZ when comparing control and *Neurod2* knockdown samples (Fig. D and E). Interestingly, in embryos electroporated with shNeurod2-1, we observed a small but significant upregulation of REELIN protein in the MZ (Fig. 5A, B and C), raising questions about the cellular source of this ectopic REELIN expression. Even though we observe a small number of neurons over-migrating into the MZ in neurons transfected with shNeurod2-1, the vast majority of neurons are localized to the CP. Whether this additional REELIN protein in the MZ is derived from the projecting dendrites of cortical neurons, or instead is a secondary response of the CR cells to reduction of *Neurod2* expression in cortical neurons is currently unclear, but this topic is worthy of future investigation.

**Figure 5.**
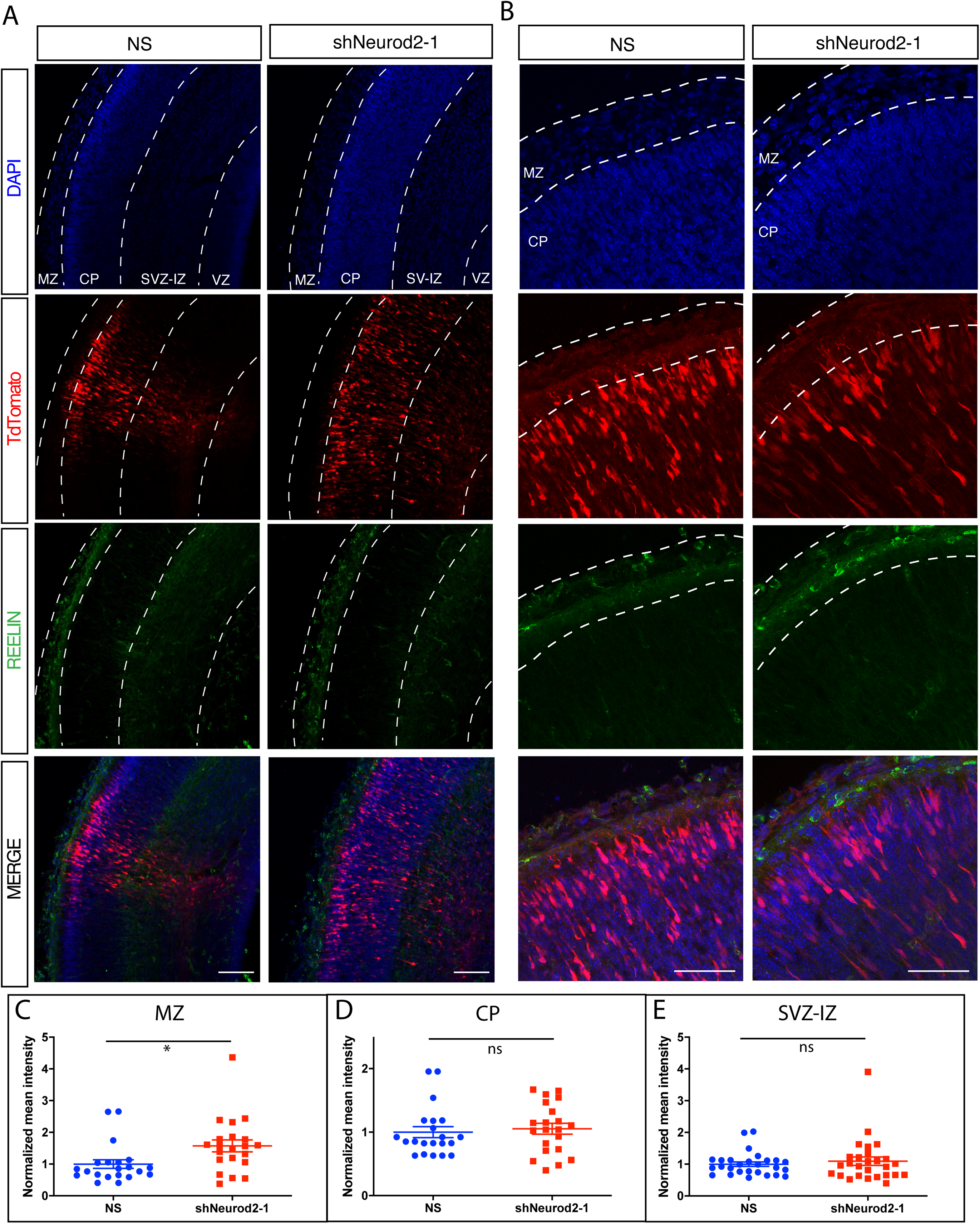
In vivo quantification of REELIN protein levels upon *Neurod2* silencing in the embryonic cortex. (A) Embryos were electroporated with shNeurod2-1 and TdTomato at E14.5, cortices were harvested at E17.5 and confocal images were acquired after REELIN immunofluorescent staining. REELIN levels were not significantly altered within the CP, SVZ-IZ and VZ. However we observed a slight but significant increase in REELIN protein with in the MZ. Scale bar : 100 µm. (B) Higher magnification images of those displayed in (A). Scale bar: 50µm. (C-E) Quantification of images displayed in (A and B). A total of 21 images acquired from n=3 embryos per group were used for quantification. *p=0.02 by unpaired two-tailed t-test. Abbreviations: MZ (marginal zone), SVZ-IZ (subventricular-intermediate zone), CP (cortical plate), NS (non-silencing), ns (non-significant).

### NEUROD2 is required for the terminal stages of cortical radial migration

Our comprehensive analysis uncovering NEUROD2 target genes functioning in neuronal migration and focused experiments on regulation of *Reln* gene expression, prompted us to test whether cortical migration is affected upon suppressing *Neurod2* expression. Toward this goal, we silenced *Neurod2* in neural progenitor cells by *in utero* transfection of shNeurod2-1 and a tdTomato marker in E14.5 embryos, then quantified the number of tdTomato-positive neurons in different cortical layers at E17.5 by confocal microscopy (Fig. 6A and B). In control samples, we observed tdTomato-positive neurons at expected locations within the ventricular zone, along the subventricular and intermediate zones, and at the cortical plate. In samples in which we silenced *Neurod2* expression, we did not observe gross defects in migration, and neurons generally appeared to migrate from the ventricular zone toward the cortical plate in a manner comparable to control samples. Specifically, multipolar migration and glial-guided migration appeared intact. However, we observed a reduction in the number of tdTomato-positive neurons in the primitive cortical zone (located right below the marginal zone), suggesting that NEUROD2 depletion leads to a defect specific to the terminal stage of migration (Fig. 6A and B). In *Neurod2* knockdown neurons, concomitant with a significant reduction of neurons localized to the primitive cortical zone, we occasionally also observed neurons which had over-migrated into the MZ (Fig. 6B and C). Taken together, our results point to a mechanism where NEUROD2 regulates the positioning of somas at the primitive cortical zone. Future experiments will clarify how NEUROD2 target genes regulate this critical step during cortical lamination.

**Figure 6.**
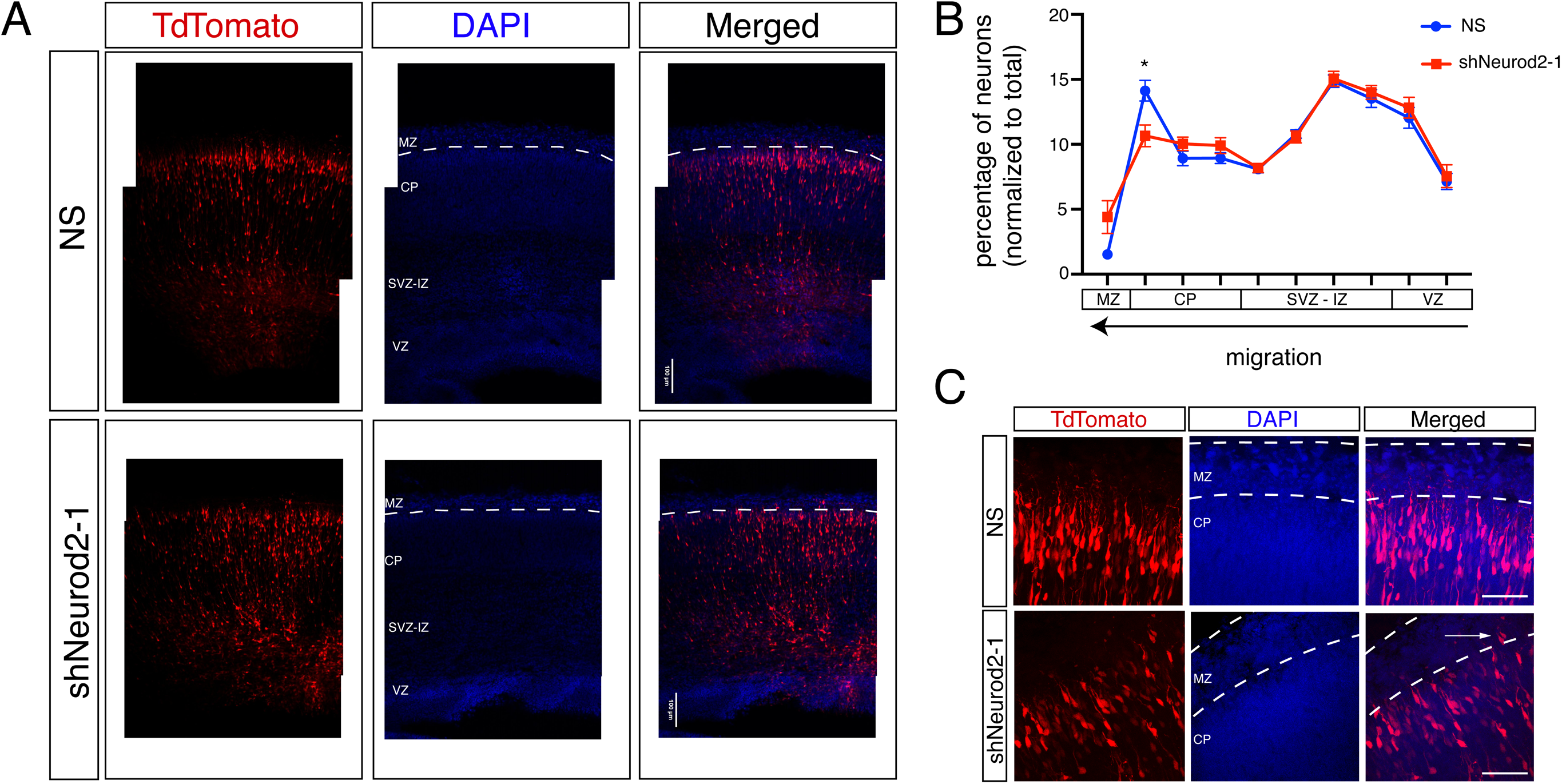
NEUROD2’s role in neuronal soma positioning to the primitive cortical zone. (A) Embryos (at E14.5) were *in utero* electroporated with tdTomato fluorescent marker and with NS (non-silencing) shRNA or with shNeurod2-1. Embryos were retrieved at E17.5, coronally sectioned and tdTomato signal was subsequently imaged by confocal microscopy. VZ (ventricular zone), SVZ-IZ (subventricular-intermediate zone), CP (cortical plate), and MZ (marginal zone) are labeled. Scale bar: 100 µm. (B) Transfected neurons (a total of n=16,184 for NS and n=19,339 for shNeurod2-1) derived from *in utero* electroporation of seven littermate embryo pairs from five independent pregnant mice were counted. Normalized percentages of tdTomato-positive neurons counted in individual zones are plotted. Significantly less neurons are localized to the upper cortical plate in neurons where *Neurod2* is knocked down. (C) Higher magnification images of the MZ and the CP in NS and shNeurod2-1 electroporated cortices. White arrow points to occasional neurons that over-migrate into the MZ of the shNeurod2-1 transfected samples. Scale bar: 50 µm. Bars represent S.E.M. *p<0.05 by two-tailed unpaired t-test.

## Discussion

Here we report a comparative analysis of the genetic targets of the TF NEUROD2 and reveal a new role for NEUROD2 in the terminal stage of cortical radial migration. We had previously reported NEUROD2 targets identified by ChIP-Seq experiments both in embryonic and postnatal cerebral cortices ^26,27^ and here we have complemented these earlier studies by a comparative analyses of these two developmental timepoints. Furthermore, in order to focus upon targets whose expression is regulated by NEUROD2, we silenced its expression in primary cortical neurons and carried out an RNA-Seq analysis. Our comprehensive and comparative analyses of these two ChIP-Seq and RNA-Seq datasets converged upon the *Reln* gene. We show that in mice, NEUROD2 binds to several different introns of *Reln* at both embryonic and postnatal stages, and this binding correlates with reduced levels of *Reln* mRNA. Even though NEUROD2 has typically been considered as a transcriptional activator, our studies point to a potentially dual function for NEUROD2; in addition to its previously reported transcription activating function ^25,42,43^, it may also act as a repressor in a context-dependent manner. In support of our findings, *Reln* expression was also previously shown to be upregulated in the embryonic cortex of *Neurod2/6* double knockout mice ^44^. However, whether this effect is a direct consequence of NEUROD2 binding to *Reln* introns or is a secondary response awaits confirmation by site-directed genome editing of these binding sites.

NEUROD2’s potential capacity to function as a transcriptional repressor agrees well with our previous results demonstrating that NEUROD2 inhibits expression of the *Stim1* gene to control calcium signaling in cortical neurons ^26^. Furthermore, several of the NEUROD2 binding peaks both in *Reln* and *Stim1* genes have also been identified in ChIP-Seq experiments carried out for the chromatin insulating factor CTCF (encodeproject.org), suggesting that NEUROD2 might be involved in establishing or maintaining higher order architecture of chromatin loops. In support of this hypothesis, NEUROD2 was recently shown to function as a major organizer of cell-type specific chromatin folding in cortical neurons, suggesting that its transcription activating or repressing functions might in fact be directly related to its role in establishing or maintaining transcriptionally isolated domains ^45^. The mechanistic details of how NEUROD2 regulates transcriptional activity in distinct genetic loci remain are of great interest during future investigations.

Past studies have suggested that NEUROD2 might have a role in regulating radial migration of excitatory cortical neurons ^27,28,46^. In addition to our current finding demonstrating NEUROD2 regulation of *Reln* mRNA expression, our past work had shown that it also binds to promoters of several genes encoding downstream signaling proteins that transduce REELIN signals, such as *Dab1* and *Fyn*, as well as the receptor gene *Lrp8* ^27^. To our surprise, upregulation of *Reln* mRNA levels upon *Neurod2* silencing does not correlate well with changes at the protein level. We observe a small but significant increase of REELIN in primary neurons, but not in embryonic sections, after *Neurod2* suppression. Immunofluorescence staining is a semi-quantitative method and might not be able to report accurately upon small changes in protein levels of REELIN. Taken together, we conclude that the major effect of NEUROD2 is most prominent at the mRNA level of *Reln*.

The control of *Reln* gene expression within a precise spatiotemporal window in the developing cortex appears to be a conserved and tightly regulated mechanism that depends upon several TFs. For example, double-knockout mice for the TFs *Pbx1/2* also exhibit ectopic *Reln* expression, leading to a dramatic inversion of cortical layering reminiscent of mouse models with defective REELIN signaling ^47^. Similarly, knockdown of the histone methyltransferase *Ezh2*, as well as deletion of *Foxg1*, result in derepression of *Reln* gene expression and are associated with radial migration and cortical development defects ^48,49^. Finally, ectopic expression of *Reln* cDNA by *in utero* electroporation into neural progenitors causes formation of neuronal aggregates in deep layers that are incapable of migrating towards the cortical plate ^50^. Taken together, these findings highlight the importance of maintaining a precise level of *Reln* gene expression for the proper development of cortical layers. Our results suggest that NEUROD2 is an additional transcriptional regulator of the *Reln* gene. Furthermore, deficiencies in NEUROD2 function might remove one layer of control against ectopic REELIN expression and potentially sensitize migrating neurons to synthetic genetic interactions with other genes functioning in REELIN signaling.

Beyond our analysis of *Reln* expression, we also generated a curated list of NEUROD2 binding targets with known roles in neuronal migration based on published literature. Interestingly, we identified a number of additional targets, including components involved in Ephrin signaling, members of the Cdk5 pathway, and additional TFs. Finally, in order to test whether NEUROD2-regulated transcription functions during radial migration of cortical neurons, we silenced *Neurod2* expression in ventricular neural progenitors by *in utero* electroporation in mice and discovered that NEUROD2 is important for proper positioning of the neuronal soma upon arrival at the primitive cortical zone. These findings suggest that NEUROD2 might impact cortical lamination by regulating multiple components of the REELIN signaling pathway or other pathways. Finally, our dataset offers novel candidate genes that may function in different aspects of cortical lamination.

## Methods

### Bioinformatic analyses of ChIP-Seq data

Previously published NEUROD2 ChIP-Seq data were used to identify differential NEUROD2 binding to genomic regions at timepoints E14.5 and P0 ^26,27^. These data were generated using three separate NEUROD2 antibodies (Abcam ab168932, ab104430 and ab109406). Histone ChIP-Seq data from E14.5 mouse embryonic cortex were produced by Bing Ren’s Laboratory, UCSD, USA ^51^ (www.encodeproject.org). ChIP-Seq peaks were visualized using Trackster ^52^ embedded within the Galaxy Project ^53^. Details of Closest Gene score calculations were previously described ^27^.

### ChIP-qPCR

Chromatin immunoprecipitation with NEUROD2 antibody was carried out using cortical tissue collected from E14.5 and P0 pups of both sexes using an antibody raised against NEUROD2 (Abcam, ab109406) and against GFP as a negative control (Santa Cruz, sc-8334). Formaldehyde-crosslinking, followed by chromatin DNA isolation from input and immunoprecipitated samples, was performed as described previously ^27^. Cq values from immunoprecipitated DNA were normalized to those from input DNA as described ^26^. Primer sequences used for qPCR experiments were as follows: Reln-peak1-F (AATGGAAACTGGCTCGCATG); Reln-peak1-R (ATCCTGAGCAATGAGTGGCT); Reln-peak2-F (TGTTTCTGTGGTCTGCTTGC); Reln-peak2-R (AAAACAATCACAGGCGAGCC); Reln-peak3-F (GCCGGACTACCCTGATGATT); Reln-peak3-R (TGGAGGAAATGGATGGCTCT); Reln-peak4-F (GGCCTCCTGTCTTACTAGCC); Reln-peak4-R (CCTGACAGATGGAGCGTTTG).

### RNA-Seq, RT-qPCR and Immunoassays

Primary cultures were prepared from E14.5 embryos as described ^26^. Cortical neuron suspensions were electroporated with shNeurod2-1, shNeurod2-2, or a non-silencing plasmid immediately before plating (Amaxa™ P3 Primary Cell 4D-Nucleofector X Kit L; Lonza, CU133 program). The generation and knockdown efficiencies of shRNA plasmids were described previously ^26^. Total RNA was isolated with Absolutely RNA Microprep Kit (Agilent Technologies) at 5 d *in vitro* (DIV). The RNA samples were further used for either RNA-seq-directed library construction or for RT-qPCR. Library construction and sequencing were carried out at Genewiz Inc. (New Jersey, USA). RNA-Seq libraries were sequenced on the Illumina HiSeq2500 platform in 50 bp single-end format. After quality filtering of reads, sequences were mapped onto the genome with TopHat ^54^, assembled onto transcripts with Cufflinks ^55^ and differential expression was determined with Cuffdiff ^55^. Raw and processed RNA-Seq data are available at Gene Expression Omnibus (http://www.ncbi.nlm.nih.gov/geo/) under the accession number GSE131494. For RT-qPCR, cDNA was synthesized with Transcriptor high-fidelity cDNA synthesis kit (Roche) and RT-qPCR was performed using Luminaris HiGreen qPCR Master Mix (Thermo Scientific) in a CFX Connect Real-Time PCR Detection System (Bio-Rad). The primer sequences were as follows: Reln-F (TTTACACTGAGGCTGGGGAG); Reln-R (TGCCACCATCTGAACTGGAT); Gapdh-F (AATGTGTCCGTCGTGGATCTGACGTGC); Gapdh-R (TTGCTGTTGAAGTCGCAGGAGACAACC). An anti-REELIN antibody (cat no. ab78540, Abcam) was used for immunoblotting and immunofluorescence staining.

### Primary cortical cultures and Sholl analysis

Primary cultures were prepared from E14.5 embryos as described ^26^. For Sholl analysis, primary cortical neurons were transfected with shNeurod2-1 ^26^ at 2 DIV with Lipofectamine 2000 transfection reagent (Invitrogen). Neurons were fixed with 4% PFA at 5 DIV, immunostained with an antibody recognizing EGFP (Aves Lab, GFP 10-20) and imaged with a Nikon 90i Eclipse confocal microscope affixed with a 60x oil objective. Dendritic development was quantified by the Sholl analysis plug-in in ImageJ software ^33,56^.

### Animals

All animal experiments were performed in accordance with the guidelines for the care and use of laboratory animals of Koç University. The study was approved by the Institutional Animal Care and Use Committee of Koç University with license number 2014-07. For *in utero* electroporation and primary neuronal cultures, timed-pregnant mice (Balb/c strain) were generated by The Koç University Experimental Animal Laboratory.

### In utero electroporation and image analysis

Overall, our *in utero* electroporation method was based upon a previously published protocol ^57^. Briefly, 14.5 day pregnant mice were anesthetized by inhalation of isoflurane. 1 µl of DNA solution (composed of 500 ng of shRNA plasmid, 500 ng of pCAGGS-IRES-tdTomato and 0.1% Fast Green FCF) was injected unilaterally into the lateral ventricular of each embryo. 0.5 pulses at 30 V with 400 ms intervals were applied with a 5 mm tweezer electrode using an electroporator (BTX, Harvard Apparatus, ECM830). Following electroporation, embryos were placed back into the uterine cavity, and the pregnant female was sutured and placed into incubation chamber for recovery. Embryos were retrieved at developmental age E17.5, followed by brain removal and placement into 4 % PFA. Embryos were coronally sectioned to 50 µm with a cryostat (Leica CM1950). Confocal images of tdTomato-positive neurons were captured with Nikon Eclipse 90i (20X objective) or Leica DMI8 (63X oil objective) and maximum intensity projection images were generated using NIS Elements AR software. A grid of ten equal bins were overlaid onto the cortical cross-section images, and the number of tdTomato-positive neurons in each bin was counted using the cell counting tool integrated into ImageJ ^56^. These results were expressed as a percentage of total number of tdTomato-positive cells. In each experiment, littermate embryos from the same pregnant mouse were electroporated with nonsilencing and shNeurod2-1, respectively.

## Acknowledgements

We thank Ali C. Taşkın, Ahmet Kocabay, Muhammed Reşat Oguzkan Yaralı and Mehmet Yücel (Koç University Animal Research Facility, Turkey) for their help in providing timed pregnant mice; Efil Bayam (Institute of Genetics and Cellular and Molecular Biology, Strasbourg, France) for assistance with the *in utero* electroporation method; Tamer Önder (Koç University School of Medicine, Turkey) for generously sharing the Lonza nucleofector equipment, and the Molecular Imaging Core Facility of Koç University Research Center for Translational Medicine (KUTTAM) for assistance with confocal microscopy.

## Author contributions

G.G. carried out ChIP-qPCR and RNA-Seq experiments (Fig. 1, 3 and 4), and primary neuronal cultures, and Sholl analysis (Fig. 2). C.A. carried out all *in utero* electroporations, confocal microscopy and data analysis, and D.A. assisted these experiments (Fig. 5 and 6). A.K., C.D.D. and N.O. contributed to data analysis (Fig.1, 3 and 4). G.I.D. conceived the hypothesis, designed research, analyzed data and wrote the paper. All authors discussed the results and reviewed the manuscript.

## Data availability

RNA-Seq data can be downloaded from Gene Expression Omnibus (http://www.ncbi.nlm.nih.gov/geo/) under the accession number GSE131494.

